# A dominant coral parasite, *Candidatus* Aquirickettsia rohweri, resists antibiotic exposure and thermal challenge below the bleaching threshold in disease-susceptible *Acropora cervicornis*

**DOI:** 10.64898/2026.07.15.738557

**Authors:** Sunni Patton, Eddie Fuques, Lauren Speare, Grace Klinges, Erinn M Muller, Rebecca L Vega Thurber

**Affiliations:** Ecology, Evolution, and Marine Biology, University of California, Santa Barbara, Santa Barbara, CA, 93106-9620, USA; School of Biological Sciences and Center for Microbial Dynamics and Infection, Georgia Institute of Technology, 950 Atlantic Drive NW, Atlanta, GA 30332, United States; Island Conservation, 630 Water St, Santa Cruz, CA 95060; Mote Marine Laboratory, 1600 Ken Thompson Pkwy, Sarasota, FL, 34236, USA

## Abstract

The critically endangered Caribbean staghorn coral *Acropora cervicornis* hosts microbiomes frequently dominated by the putatively parasitic intracellular bacterium *Candidatus* Aquirickettsia rohweri, which is associated with reduced coral growth and heightened disease susceptibility. Whether this dominance can be disrupted through antibiotic treatment and a sequential disturbance of thermal stress, remains unknown. In this study, we exposed disease-susceptible *A. cervicornis* fragments to broad-spectrum antibiotics, sub-bleaching thermal stress, or the combination of an antibiotic pre-treatment followed by thermal stress, and tracked changes in microbiome composition and diversity across all experimental phases using 16S rRNA amplicon sequencing and quantitative PCR (qPCR). We find that while the minor microbial fraction exhibits sustained compositional shifts in response to treatment, *Ca.* Aquirickettsia rohweri is resilient to antibiotic and thermal perturbation and may in fact increase in abundance following antibiotic exposure, suggesting that its dominance is actively maintained and not readily displaced by current disease mitigation strategies.These results indicate that antibiotic intervention is unlikely to be a viable strategy for disrupting *Ca.* A. rohweri dominance in disease-susceptible *A. cervicornis*, underscoring the urgency of understanding its transmission routes to inform microbiome rescue efforts.

## Introduction

Host-associated microbiomes are often regarded as integral to proper host development and sustained health, as many bacteria complement host fitness, extend metabolic capacity, and confer protection against pathogens (Lynch & Hsiao, 2019; McFall-Ngai et al., 2013; Shabat et al., 2016; Sommer & Bäckhed, 2013). Microbiome-mediated immune protection can occur through physical occlusion and competitive exclusion of pathogens (Endt et al., 2010), as well as through the production of antimicrobial compounds and nutrient blocking that limit pathogen proliferation (Spragge et al., 2023). In many systems, microbiomes that exhibit high taxonomic and functional diversity, functional redundancy, and non-dominance by any single taxon are often associated with coordinated ecosystem function and an increased capacity for hosts to appropriately respond to, or resist, disturbances (Lozupone et al., 2012; Yachi & Loreau, 1999). However, defining a “healthy” microbiome structure is highly context-dependent, as microbiome composition is continuously shaped by host genetics, environmental signals (Glasl et al., 2019), and microbe-microbe interactions (Lozupone et al., 2012; Ritchie, 2006; Zaneveld et al., 2017).

Scleractinian corals rely heavily on their associated bacterial communities for nutrient cycling (Kimes et al., 2010; Rädecker et al., 2015), pathogen exclusion, and tolerance to environmental perturbations (Cárdenas et al., 2022; Marangon et al., 2021; Reshef et al., 2006), and often harbor complex and discrete microbiomes within each body site microhabitat (Sweet et al., 2011a). In addition to structural complexity, coral-associated microbial communities are drastically restructured throughout ontogeny, differ by environment, and often differ both across and within host species (Bernasconi et al., 2019; Ricci et al., 2022). The degree to which microbiome composition is conserved within a host species reflects the interplay between host filtering mechanisms and local environmental conditions, and there is considerable within-species variation in microbiome structure (Rosales et al., 2019; Williams et al., 2022). Understanding what maintains particular microbiome configurations, and whether those configurations are beneficial or maladaptive, remains an open question in coral biology.

The critically endangered Caribbean staghorn coral, *Acropora cervicornis*, provides a compelling system for investigating the relationship between microbiome structure and host health. Florida populations of *A. cervicornis* display few genetic variants with resistance to White Band Disease (WBD), with the majority of genotypes exhibiting high disease susceptibility (Klinges et al., 2020; Muller et al., 2018; Williams et al., 2022). While host genetic factors likely contribute to this susceptibility (Vollmer et al., 2023), disease-susceptible genotypes are also characterized by a distinctive microbiome structure: overwhelming dominance by a single bacterial taxon, *Candidatus* Aquirickettsia rohweri (Klinges et al., 2020; Williams et al., 2022). This intracellular, putatively parasitic bacterium is incapable of synthesizing its own ATP and must exploit the host by siphoning ATP via an ATP/ADP translocase (Klinges et al., 2019). Although corals harboring *Ca.* A. rohweri appear outwardly healthy under ambient conditions, its presence is associated with reduced coral growth rates (Shaver et al., 2017), and under nutrient enrichment, *Ca.* A. rohweri proliferates and upregulates several genes involved in metabolism, nutrient uptake, and host interaction and virulence (Speare et al., 2026). Importantly, when *Ca.* A. rohweri is lost, as occurs under thermal stress conditions and *Vibrio coraliilyticus*-mediated bleaching, the vacated niche space may become available to opportunistic bacteria, thereby worsening bleaching and disease outcomes (Klinges et al., 2020; Speare et al., 2025). Furthermore, despite reliance on the coral host for ATP and amino acids, *Ca.* Aquirickettsia rohweri is not vertically transmitted and instead must be acquired from the environment at some point during coral development (Baker et al., 2022). The absence of vertical transmission further distinguishes this parasite and raises pressing questions about how it is acquired, established, and maintained at such high relative abundances, and whether its dominance is amenable to intervention.

Antibiotic treatment represents one potential avenue for disrupting bacterial dominance within host microbiomes, as broad-spectrum antibiotics can deliberately deplete abundant taxa and restructure community composition, often without obvious negative effects on overall host health (Neely et al., 2020; Patton et al., 2025; Sweet et al., 2014). However, the degree to which antibiotic-mediated reductions in dominant members persist, and how the broader community responds, is a facet of coral microbiome research actively being studied (Bent et al., 2021; Dunphy et al., 2021; Patton et al., 2025). *Ca.* Aquirickettsia rohweri is highly sensitive to thermal stress, yet whether combining antibiotic exposure with subsequent thermal stress might reduce *Ca.* A. rohweri burden is unknown. Therefore, in this study we exposed disease-susceptible *A. cervicornis* from Florida to antibiotics, elevated temperature, or both, and tracked microbiome compositional changes and *Ca.* Aquirickettsia rohweri absolute abundance over time and treatment.

## Experimental Procedures

### Experimental design and sample collection

In March 2022, 144 White Band Disease (WBD) susceptible *Acropora cervicornis* (genotype ML-50) fragments were collected from Mote Marine Laboratory’s *ex-situ* nursery. Fragments acclimated to outdoor raceway conditions for one week in Mote’s Climate and Acidification Ocean Simulator (CAOS) system. After acclimation, coral fragments were then randomly assigned one of four treatments: ‘no treatment’, ‘antibiotics’, ‘temperature’, and ‘antibiotics + temperature’ and placed into the corresponding 5-gallon, non-flow through tanks containing 6 L of water and an aquarium pump. Each treatment had four replicate tanks with nine coral fragments in each tank. Tanks were randomly assigned a position in the raceways, with 8 tanks per raceway. The biphasic experiment began with a 96-hour antibiotic challenge, followed immediately by a thermal stress challenge. A mixture of ampicillin, streptomycin, and ciprofloxacin was used for the antibiotic challenge, such that when added to the tanks, ampicillin and streptomycin would each be at a final concentration of 10 mg/L and ciprofloxacin 1 mg/L. These antibiotics at these doses have been used in several other studies (Connelly et al., 2022, 2023; Gilbert et al., 2012; Sweet et al., 2014), and do not result in obvious host phenotypic stress. For a detailed protocol, see Patton et al., 2025 (Patton et al., 2025).

Every 24 hours during the antibiotic exposure, half of the water volume was removed, replenished, and re-dosed with antibiotics to maintain a consistent concentration. Entire coral fragments were collected prior to the antibiotic challenge (T0), midway through (T2), and at the end of the antibiotic challenge before beginning the temperature ramp (T4). All samples were placed into 7 mL of DNA/RNA Shield (Zymo Research, Irvine, CA, USA) and stored at −80 C until further processing. Immediately after the 96-hour antibiotic challenge, antibiotic-exposed tanks were replaced with clean tanks in order to safely allow for a flow-through tank design for the remainder of the experiment. We then began a temperature ramp from 27.5 C to 30 C by 0.5 C/day, after which the temperature was held for the remainder of the experiment. Incoming water from the header tank was set at a consistent 27.5 C to reduce system stress, while the water in the raceway was heated to the set temperature to gradually increase the internal tank temperature. Our ramp period was 10 days, and the corals were at peak temperature for a total of 20 days. After beginning the temperature challenge, samples were collected every five days (until T34) as previously described. The experimental design can be seen in Figure 1.

**Figure 1.**
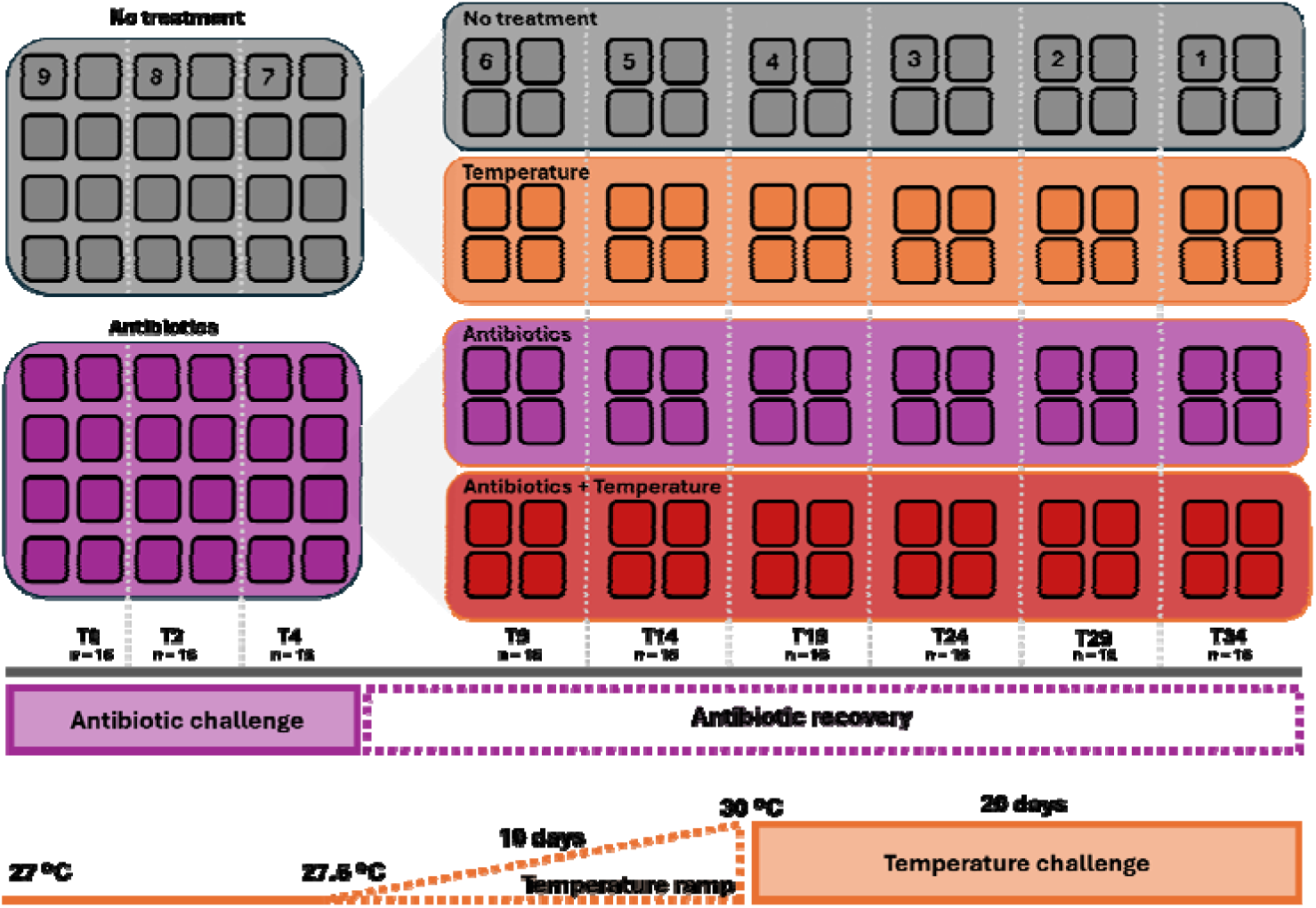
Experimental design displaying the treatments, sample sizes, and phases of the experiment. Four tanks per treatment (‘no treatment’, ‘antibiotics’, ‘temperature’, and ‘antibiotics + temperature’) were used. Individual coral frags (n = 144) were collected at each time point from each of the 16 tanks (n = 16 per timepoint). The numbers within the boxes represent the number of frags remaining in each tank prior to the sampling point. All tanks within a timepoint have the same number of frags. Prior to the experiment, the 144 frags were split into ‘no treatment’ and ‘antibiotics’. The antibiotic group (n = 72) were exposed to antibiotics for four days and samples were collected at T0, T2, and T4. After the antibiotic treatment, both the ‘antibiotics’ and ‘no treatment’ groups were further evenly split into ‘temperature’ and ‘antibiotics + temperature’ (n = 36). Following the antibiotic challenge, the water temperature in the temperature treatment groups was increased at 0.5□/day (10 days). In the CAOS system, the water temperature was ultimately set to 32□but the internal tank temperature only reached ∼30□. Therefore, there was a 10-day ramp period to reach 30□. After the ramp, the tanks were held at 30□for the remainder of the experiment (20 days). Given that antibiotics were not administered after the T4 timepoint, the remainder of the experiment for the antibiotic-treated groups served as an antibiotic recovery period.

### Sample DNA extraction, 16S rRNA gene library preparation, and sequencing

Coral samples were thawed and approximately 5 polyps were removed using sterile bone cutters. Polyps were placed into DNA extraction tubes along with 100 µL of DNA/RNA Shield that the fragment was stored in. Sample DNA was then extracted using the DNeasy PowerSoil Pro Kit (Qiagen) according to the manufacturer’s instructions. The V4 region of the 16S rRNA gene was amplified via one-step polymerase chain reaction (PCR). 25 µL PCR reactions were made using 10 µL of Platinum II *Taq* Hot-Start PCR Master Mix (2x) (Invitrogen), 2.5 µL each of 10 µM primers 515F (5’ - GTGYCAGCMGCCGCGGTAA - 3’) (Parada et al., 2016) and 806R (5’ -GGACTACNVGGGTWTCTAAT -□3’) (Apprill et al., 2015) with attached barcodes. Four negative controls (MilliQ water) were sequenced alongside the samples. The template DNA was amplified using the following thermocycler parameters: initial denaturation at 94 C for 2 min, followed by 35 cycles of denaturation at 94 C for 30 s, annealing at 60 C for 30 s, and extension at 68 C for 60 s, followed by a single final extension step at 68 C for 10 min. Amplified PCR products were then purified using Agencourt AMPure XP beads (Beckman Coulter) following the manufacturer’s guidelines. However, 80% ethanol was used for the washing steps rather than the recommended 70%, and a 5-minute drying step was included after the final ethanol wash to evaporate excess ethanol. After purification, DNA concentrations were quantified using the BioTek Synergy H1 multi-mode plate reader. Libraries were pooled at equimolar concentrations before paired-end 2□×□300 bp sequencing using the Illumina MiSeq at Oregon State University’s Center for Quantitative Life Sciences (CQLS).

### Quantitative PCR (qPCR)

Quantitative PCR was performed on all coral samples in triplicate targeting both the *tlc1* gene to identify *Ca.* A. rohweri and the actin gene as an endogenous control (Wright et al., 2019) following a method adapted from Klinges et al., 2022 (Klinges et al., 2022). In each reaction for each primer set, master mixes were created using 10 µL of iQSybr (BIO-RAD), 0.6 µL of 0.3 µM forward and reverse primers, and 8.3 µL of water. In each well, 0.5 µL of 20 ng/µL template DNA was used. In each plate, DNA from a ML-50 sample and an *A. hyacinthus* sample was used as interplate calibrators (IPCs) for the *tlc1* and actin genes, respectively. Six negative controls, three for each gene target, were also run in triplicate in each plate. Plates were run on the BIO-RAD 96XTouch using the same PCR parameters as previously described. Data for each gene target were first separated and the thresholds for each plate were adjusted such that the Cq values of the IPCs were within 0.25 standard deviations. Following threshold adjustment, triplicate values were averaged and the Cq value of the actin gene was subtracted from the Cq value of the *tlc1* to obtain ΔCt for each sample. To determine relative quantification of *Ca.* A. rohweri abundance, the no treatment samples were set as the reference to which the experimental groups were compared to within each timepoint. The mean ΔCt of the experimental treatment group was then subtracted from the mean ΔCt of the reference group to obtain the ΔΔCt value. Relative quantification was then determined by 2^-ΔΔCt^. Significant differences in *Ca.* A. rohweri abundance were determined using the Kruskal-Wallis and Dunn post-hoc tests.

### Microbiome data preprocessing

A total of 148 samples were demultiplexed by the CQLS, and individual forward and reverse quality profiles were generated using FastQC and MultiQC (Ewels et al., 2016). Using a two-step cutadapt (Martin, 2011) approach to remove forward, reverse, and reverse complements, all remaining primer sequences were removed. Reads were then imported into RStudio (v4.4.2), after which the standard DADA2 (Callahan et al., 2016) was followed. In brief, forward and reverse reads were truncated at 250 and 200 bp, respectively, leaving 12,608,343 reads remaining (5.8% of reads lost). Given the large dataset size, the number of bases used to calculate error was set at 5×10^8^ (five times more than the default value), as it provided a middle ground between utilizing more data for calculating the error while still minimizing the computational effort. Denoised forward and reverse sequences were then merged into contigs, and only those within the expected amplicon size range were retained for analysis (250 - 256 bp). Chimeric, chloroplast and mitochondrial sequences were subsequently removed along with any amplicon sequence variants (ASVs) not annotated beyond the kingdom level – resulting in a total of 12,071,891 reads and 4,749 taxa remaining. After building the first phyloseq object (McMurdie & Holmes, 2013), contaminant ASVs were identified and removed (29 ASVs) via the *decontam* package (Davis et al., 2018) using the combined detection method at a 0.25 threshold, which utilizes both DNA quantification values and prevalence data within negative controls. Negative controls were then removed, leaving a total of 144 samples, 4,709 ASVs, and 12,065,895 reads.

### Microbiome and statistical analysis

Prior to calculating microbiome diversity metrics, ASVs that had both low frequency (present in only one sample) and low prevalence (reads below the first quartile) were pruned from the phyloseq object, as these do not likely represent biologically relevant taxa. A phylogenetic tree was then created by aligning the sequences using MAFFT (Katoh & Standley, 2013), followed by tree construction using IQ-TREE 2 (Minh et al., 2020) which was rooted at the midpoint prior to analyses. The phyloseq object was then rarefied such that all samples had 52,686 reads (across 3,228 ASVs; Figure S1). To visualize community composition, taxa counts were transformed into relative abundances and the top 15 most abundant ASVs were identified, and replicate samples from each treatment group were then merged to display mean relative abundance (Figure 2). The pruned, rarefied phyloseq object was then used to calculate Shannon and Faith’s Phylogenetic Diversity. To identify significant drivers of alpha diversity, generalized linear models were used, as the data follow a gamma distribution. Data were subset by experimental phase (antibiotic challenge, temperature ramp, and temperature challenge), treatment and time were set as fixed effects, and tank was set as the random effect for each model. Pairwise comparisons between treatment groups within each experimental phase was assessed using the emmeans package (Lenth, 2016), and the p-values were adjusted using the Tukey adjustment.

**Figure 2.**
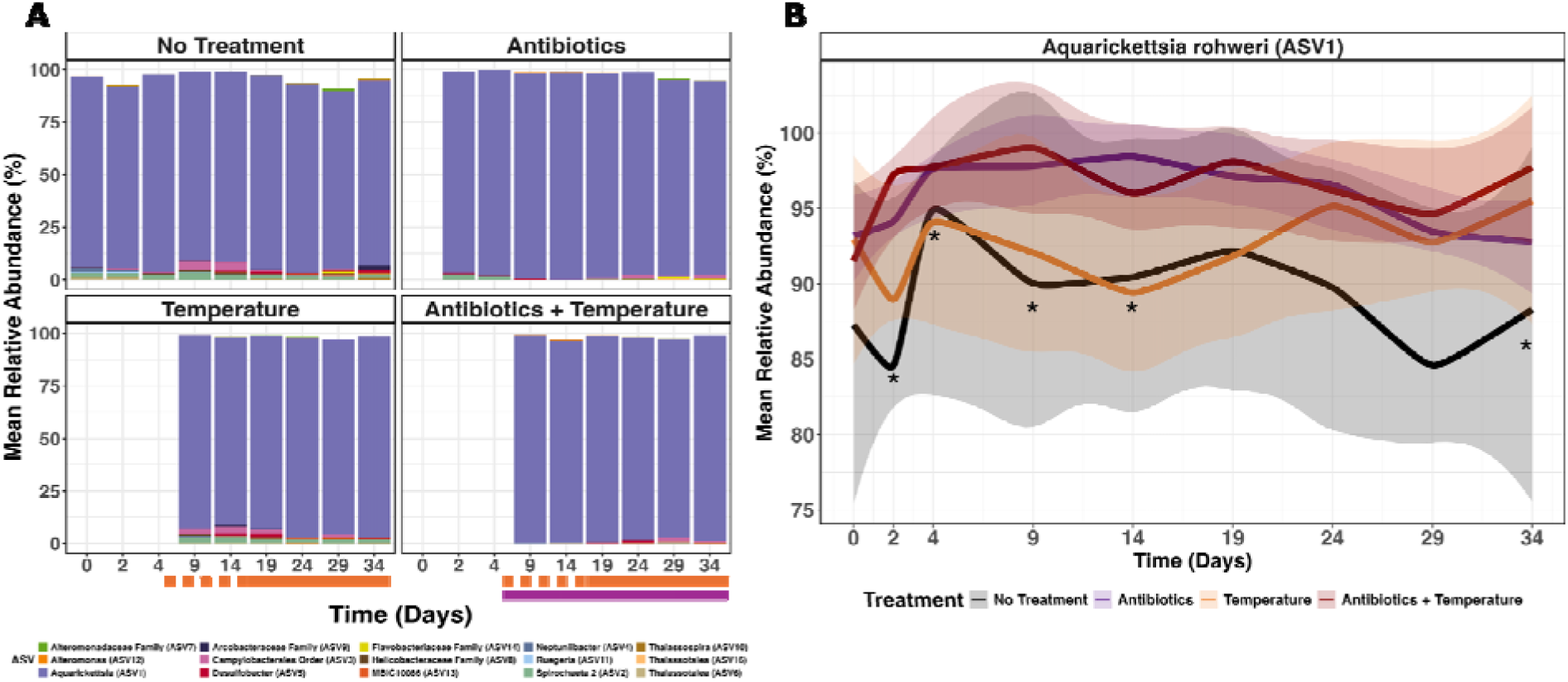
A) Mean relative abundance of the top 15 most abundant ASVs. Each bar represents the combined samples within a time point. The T0 bar in the ‘no treatment’ group includes 16 combined samples, the T2 and T4 bars in the ‘no treatment’ group each include 8 combined samples, and the T2 and T4 bars in the ‘antibiotics’ and the ‘antibiotics + temperature’ treatment groups each contain 4 combined samples. All other bars include 4 combined samples. The days represent the number of days after the beginning of the experiment. Time points without bars do not indicate missing data. Instead, those samples were added to either the ‘no treatment’ group or the ‘antibiotics’ group, as the temperature phase had not yet started. The dashed and solid orange lines represent the temperature ramp and peak temperature challenge, respectively. The purple line represents antibiotic recovery. B) Mean relative abundance of *Ca.* A. rohweri over time. Stars indicate a significant difference in relative abundance in a treatment group compared to the ‘no treatment’ group. Statistical significance was determined using raw relative abundance values (not the mean values displayed in the plot) and can be seen in Supporting Information File 1.

For beta diversity analyses, data from the non-rarefied phyloseq object were subset by experimental phase and further centered-log ratio (CLR) transformed due to the compositional and sparse nature of sequencing data (Martino et al., 2019). For each subset, Aitchison distance was calculated for each sample and data were oriented using a Principal Component Analysis (PCA). Unweighted Unifrac distance was calculated using data from the non-rarefied, non-transformed phyloseq object and ordinated using a PCoA. For both metrics, significance was determined via PERMANOVA using *adonis2* in the *vegan* package (Oksanen et al., 2025; v2.6.10) with treatment and time set as fixed effects. Pairwise interactions were calculated by subsetting the distance matrix and metadata and repeating *adonis2*. Data were subset after calculating the distance matrix in order to preserve the distance values of the community. Beta dispersion (as distance to centroid) was determined using Euclidean distances that were calculated from the CLR-transformed data and from the unweighted Unifrac distances. Statistically significant differences were determined using the *betadisper* and *permutest* functions in the *vegan* package (Oksanen et al., 2025; v2.6.10) with fdr p-value correction.

Differential abundance analysis was performed using MaAsLin2 (Mallick et al., 2021). Samples were subset by experimental phase, treatment and time were set as fixed effects, and tank ID was set as the random effect. In each phase subset, each experimental treatment group was compared against the phase control samples. These samples included T2 and T4 ‘no treatment’ samples, T9 and T14 ‘no treatment’ samples, and T19, T24, T29, and T34 ‘no treatment’ samples within the antibiotic challenge, temperature ramp, and temperature challenge phases, respectively. In the MaAsLin2 formula, data were also CLR-normalized. Therefore, the output coefficient represents the CLR-normalized abundance relative to the reference.

## Results

### Parasitic *Ca*. Aquirickettsia rohweri is not negatively affected by antibiotics, sub-bleaching thermal stress, or a sequential disturbance of the two

*Ca.* Aquirickettsia rohweri (ASV1) was identified as the most dominant ASV in every sample (Figure 2, Figure S2), ranging in relative abundance from 76.3% to 99.8% in the ‘no treatment’ and ‘antibiotics’ groups, respectively. Additionally, *Ca.* A. rohweri relative abundance was not negatively affected by antibiotic treatment, sub-bleaching thermal stress, or by thermal stress immediately following an antibiotic challenge (Figure 2A). In fact, *Ca.* A. rohweri had significantly higher relative abundance in antibiotic-treated samples (T2: 95.5% ± 1.63%, T4: 97.7% ± 0.45%) compared to untreated samples (T2: 85.4% ± 4.07%, T4: 93.7% ± 1.55%) (Figure 2B; p = 0.002 and p = 0.001 for T2 and T4, respectively; Supporting Information File 1). At T9, the ‘antibiotics + temperature’ samples had significantly higher *Ca.* A. rohweri relative abundances (99% ± 0.78%) compared to the ‘no treatment’ (89.2% ± 1.74%) (p = 0.012) and ‘temperature’ groups (94.7% ± 1.92%) (p = 0.047). However, further into the temperature ramp at T14, only the ‘antibiotic’ group displayed significantly higher *Ca.* A. rohweri relative abundances (98.4% ± 1.81%) compared to the ‘temperature’ samples (91% ± 1.73%) (p = 0.023). At the final timepoint (T34) only the ‘antibiotics + temperature’ group displayed significantly higher relative abundance of *Ca.* A. rohweri (98.2% ± 0.78%) compared to the ‘no treatment’ group (90.3% ± 4%) (p = 0.036).

Furthermore, when samples were grouped by phase (antibiotic challenge, temperature ramp, or temperature challenge) rather than individual timepoints, differential abundance analysis revealed that *Ca.* A. rohweri relative abundance was significantly higher in the antibiotic-treated samples compared to the controls during the antibiotic challenge phase (Figure 3A). It must be noted, however, that time was also a significant driver of increased *Ca.* A. rohweri relative abundance during the antibiotic phase – yet it was not a driver of changes in *Ca.* A. rohweri relative abundance in any other phase (Supporting Information File 1). During the temperature challenge phase, *Ca.* A. rohweri relative abundance was also significantly higher in all treatment groups (antibiotics only, temperature only, and antibiotics + temperature) compared to the combined control samples from T19, T24, T29, and T34 (Figure 3A), yet *Ca.* A. rohweri relative abundance was not significantly different from the controls during the temperature ramp phase.

**Figure 3.**
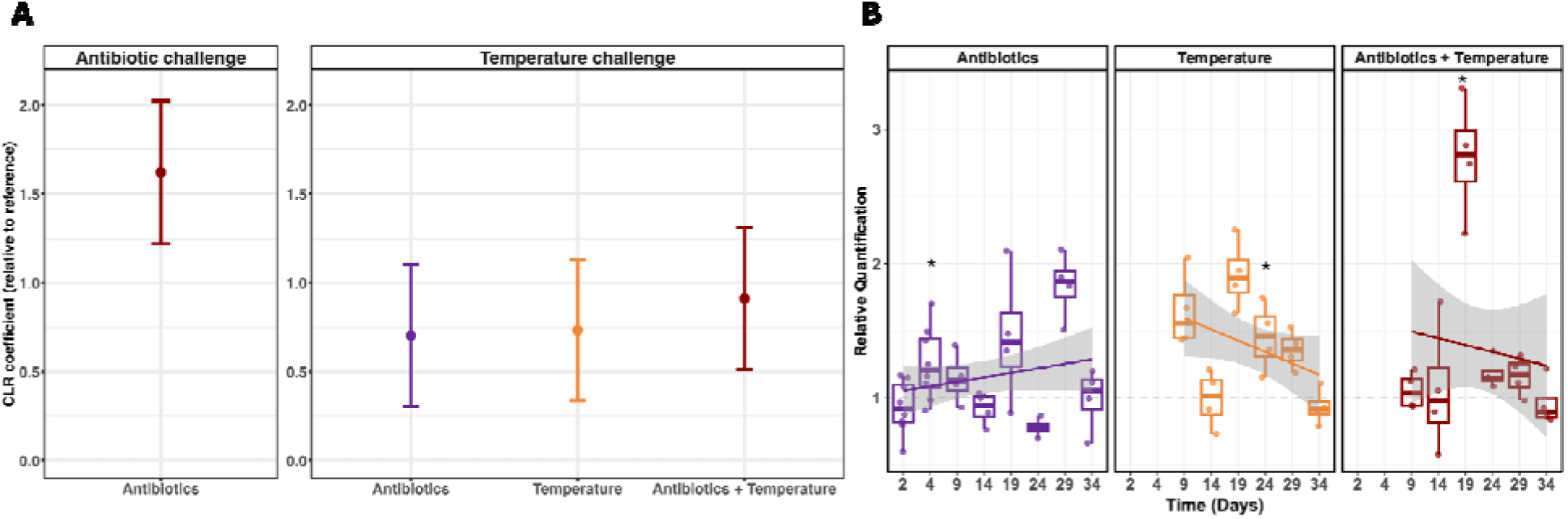
A) *Candidatus* Aquirickettsia rohweri differential abundance, as determined by MaAsLin2. Within each experimental phase (antibiotic challenge and temperature challenge), the treatment samples are compared against the ‘no treatment’ samples within the specific phase. The coefficient value represents the direction of change, as well as the CLR-normalized abundance relative to the reference. B) Boxplot of *Ca.* A. rohweri relative quantification as determined by qPCR. Values from each timepoint are relative to the ‘no treatment’ group within that same timepoint. The line represents the conditional mean of the data, and the shaded region represents the 95% confidence interval.

Due to the compositional nature of sequencing data, changes in relative abundance may not accurately capture changes in true abundance. Therefore, we also utilized qPCR to explore changes in *Ca.* A. rohweri total abundance. The above general trends are supported by the qPCR data which revealed that the antibiotic-treated samples at T4 (antibiotic challenge phase) had significantly higher *Ca.* A. rohweri abundance compared to control samples at the same time (Figure 3B). At T19 and T24, the temperature-only samples and antibiotic + temperature samples displayed significantly higher *Ca.* A. rohweri abundances relative to the control samples at the same time (Figure 3B). Despite individual timepoints displaying higher *Ca.* A. rohweri abundances in each of the treatment groups, only the ‘antibiotics’ group follows a positive trajectory of conditional means, whereas both the ‘temperature’ and ‘antibiotics + temperature’ groups’ conditional means follow a negative slope (Figure 3B).

### Antibiotics drive sustained changes in microbiome community structure and diversity within the rare biosphere

Overall Shannon diversity in this disease-susceptible *A. cervicornis* genotype is remarkably low (Figure 4), largely influenced by the dominance of a single *Ca.* Aquirickettsia rohweri ASV (Figure 2A). Despite this low diversity, however, antibiotic challenge resulted in a sustained reduction in Shannon diversity compared to the ‘no treatment’ controls (Figure 4). During the antibiotic challenge phase (T0 - T4), the antibiotic-treated group had significantly lower Shannon and Faith’s Phylogenetic Diversity compared to the ‘no treatment’ group (Figure 4C; p < 0.0001 and Figure 4D; p < 0.0001, respectively). During the temperature ramp (T9 and T14), the ‘antibiotics’ and ‘antibiotics + temperature’ treatment groups displayed lower Shannon diversity than both the ‘no treatment’ (p < 0.0001 and p = 0.0004, respectively) and ‘temperature’ groups (Figure 4C; p < 0.0001 and p = 0.0003, respectively), while the ‘no treatment’ and ‘temperature’ groups did not significantly differ. Additionally, Shannon diversity was not significantly different between the ‘antibiotics’ and ‘antibiotics + temperature’ groups. Conversely, Faith’s PD was not found to significantly differ between any of the groups during the temperature ramp phase. During the temperature challenge (T19 - T34), all experimental treatment groups displayed lower Shannon diversity compared to the ‘no treatment’ group (Figure 4C; ‘antibiotics’ p = 0.0002, ‘temperature’ p = 0.0073, ‘antibiotics + temperature’ p < 0.0001). Interestingly, Shannon diversity for the ‘temperature’ treatment was significantly higher than the ‘antibiotics + temperature’ group (p = 0.009), yet was not significantly different from the ‘antibiotics’ group (Figure 4C).

**Figure 4.**
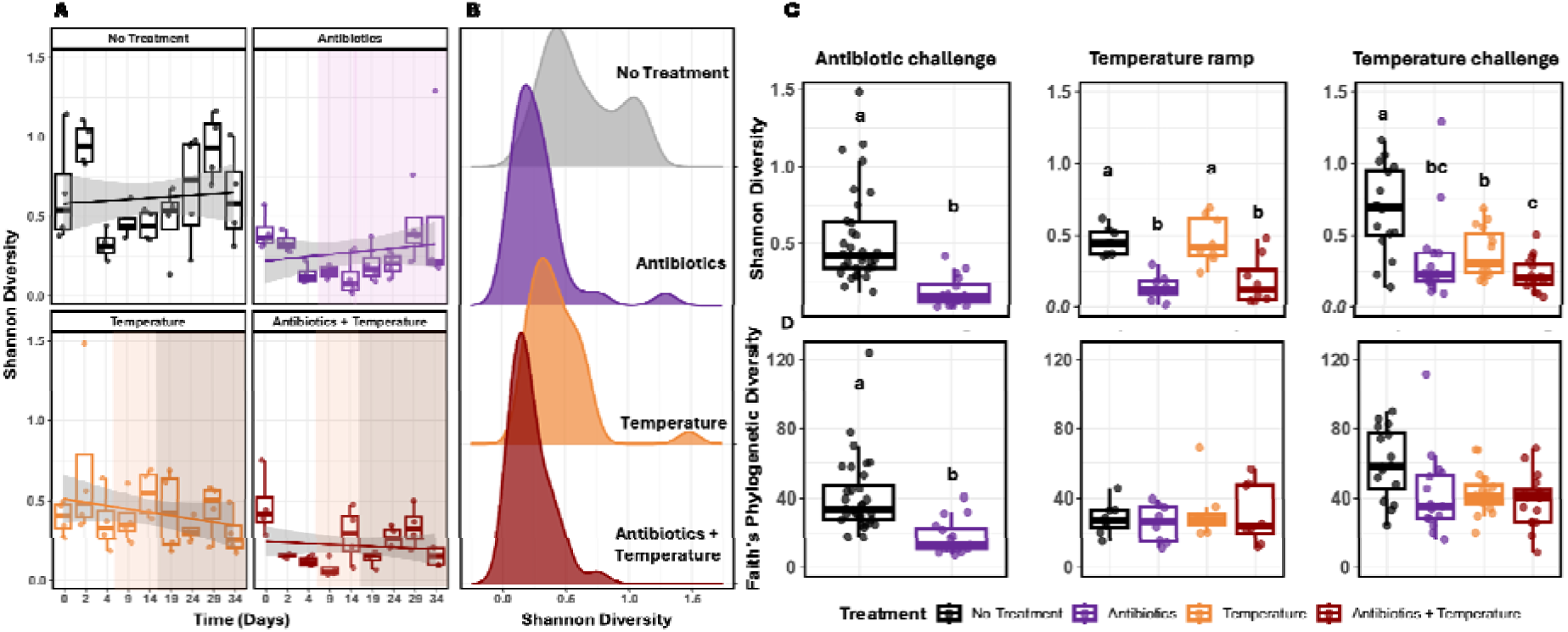
A) Shannon Diversity within each treatment group over time. The light orange overlay highlights the samples that were taken during the temperature ramp phase and the dark orange overlay highlights the samples that were taken during the temperature challenge phase. Note that the light and dark overlays in the ‘antibiotics + temperature’ group also represents the antibiotic recovery period. The grey ribbon represents the 95% confidence interval. B) Ridgeline plot displaying overall Shannon Diversity by treatment group. C) Comparative boxplots displaying Shannon Diversity between treatment groups within the three experimental phases. Boxplots not sharing a letter are statistically significantly different from one another. D) Boxplots displaying Faith’s Phylogenetic Diversity between treatment groups in each of the three experimental phases.

Furthermore, beta diversity analyses revealed that during the antibiotic challenge, community composition was significantly different between the ‘no treatment’ and ‘antibiotics’ groups for both Aitchison distance and unweighted Unifrac (Figure 5A and 5C; p = 0.0002). The effects of time were also found to drive significant differences in beta diversity, yet after accounting for the effects of time in the permutation test, treatment remained a significant driver of microbiome compositional differences. Immediate compositional shifts were seen for both distance metrics during the antibiotic challenge phase (Figure 5A and 5C) These trends continued throughout the temperature ramp and temperature challenge phases. In the temperature ramp phase, both of the antibiotic-treated groups displayed significantly different community composition compared to the no treatment and temperature only groups for both Aitchison and Unweighted Unifrac (Figure 5A and 5C; Supporting Information File 1). No significant differences were found between ‘no treatment’ versus ‘temperature’ and ‘antibiotics’ versus ‘antibiotics + temperature’ in the temperature ramp phase. During the temperature challenge, sustained differences in community composition were observed throughout the temperature challenge in which the ‘antibiotics’ and ‘antibiotics + temperature’ groups displayed significantly different microbiome compositions to the ‘no treatment’ group by both distance metrics (Figure 5A and 5C; Supporting Information File 1). Beta diversity was also significantly different between the antibiotic-treated groups compared to the ‘temperature’ group for Aitchison distance (Figure 5A; Supporting Information File 1), while only the ‘antibiotic + temperature’ samples were significantly different from ‘temperature’ group for Unweighted Unifrac (Figure 5C; Supporting Information File 1). Furthermore, we observed significant shifts in community composition in the ‘temperature’ group compared to the ‘no treatment’ group during the temperature challenge for both beta diversity metrics (Figure 5A and 5C; Supporting Information File 1). Beta dispersion, as distance to centroid, was significantly lower in the antibiotic-treated group compared to the control group during the antibiotic challenge phase, yet only for Aitchison distances (Figure 5B, p = 0.001). During the temperature challenge phase, both ‘temperature’ and ‘antibiotics + temperature’ had significantly lower dispersion than the control group for Aitchison distances (Figure 5B), whereas the ‘antibiotics’ and ‘antibiotics + temperature’ groups displayed higher dispersion compared to the ‘temperature’ group for Unweighted Unifrac (Figure 5D).

**Figure 5.**
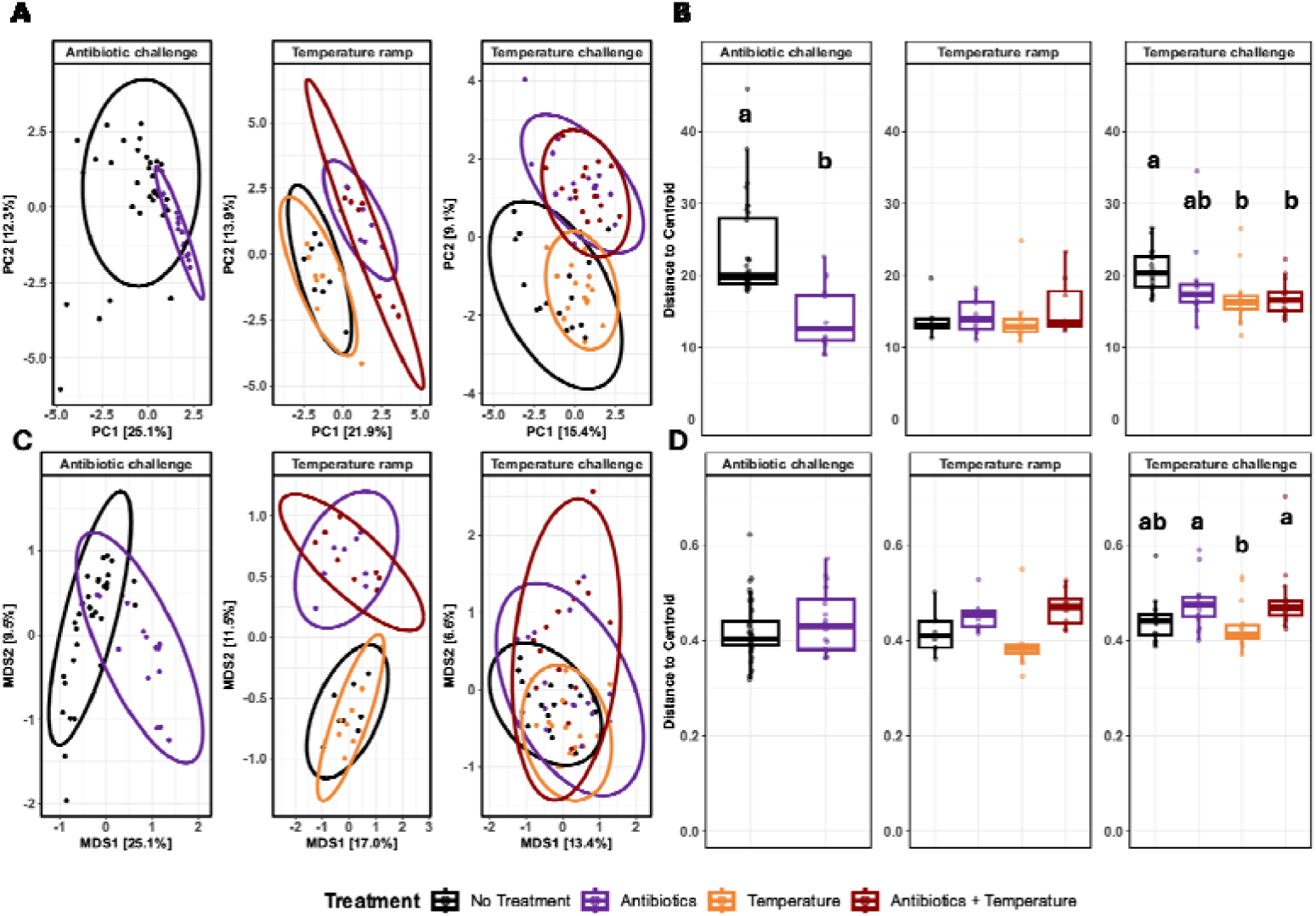
A) PCA displaying beta diversity for CLR-transformed Aitchison distances and B) a boxplot displaying beta dispersion as distance to centroid. C) PCoA displaying beta diversity for unweighted Unifrac distances and D) a boxplot displaying beta dispersion as distance to centroid. Due to the overwhelming dominance of *Ca.* A. rohweri, observable shifts in microbiome composition are difficult to identify (Figure 2), yet both beta diversity and differential abundance analyses revealed that antibiotic challenge results in sustained effects on the rare fraction of the microbiome (Figure 5 and Figure 6).

Upon removing the dominant *Ca.* A. rohweri ASV1 from the analysis, the abundance patterns of rarer taxa become more clear (Figure 7). After *Ca.* A. rohweri, the second most abundant ASV was *Spirochaeta 2* (ASV2), which ranged from 0% and 4.5% relative abundance, and was observed in higher abundance in the ‘no treatment’ group throughout the duration of the experiment, during active antibiotic treatment (T2 and T4), and throughout the temperature-only group (Figure 7A). During the antibiotic recovery phase for the ‘antibiotics’ and ‘antibiotics + temperature’ group, however, *Spirochaeta 2* (ASV2) was significantly reduced in relative abundance (Figure 6 and Figure 7). During the temperature ramp phase, the Campylobacterales Order (ASV3) had significantly higher relative abundance in the ‘no treatment’ and ‘temperature’ groups compared to the antibiotic-treated groups (Figure 6B and Figure 7A), yet this ASV ultimately increased in relative abundance during the temperature challenge phase to levels comparable to the ‘no treatment’ and ‘temperature’ groups (Figure 7A and 7B). Helicobacteraceae Family (ASV8) was found at significantly lower relative abundance in the temperature ramp phase in the antibiotic-treated groups compared to the ‘no treatment’ and ‘temperature’ groups (Figure 7B).

**Figure 6.**
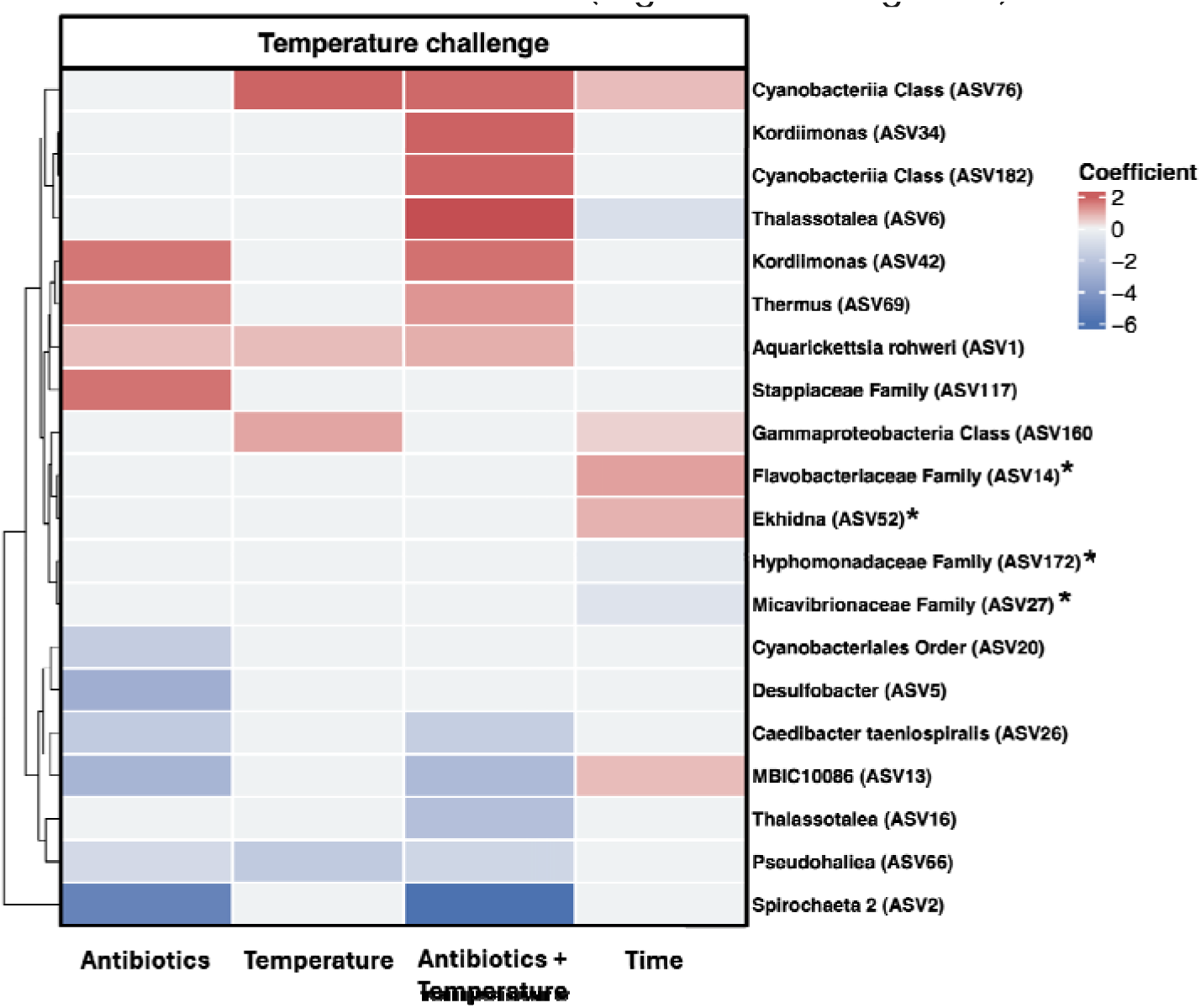
Heatmap of differentially abundant taxa during the temperature challenge phase, as determined by MaAsLin2. Each treatment group within the temperature challenge phase was compared against the ‘no treatment’ samples within the phase. The coefficient value represents the direction of change, as well as the CLR-normalized abundance relative to the reference. The stars denote ASVs that were differentially abundant only over time.

**Figure 7.**
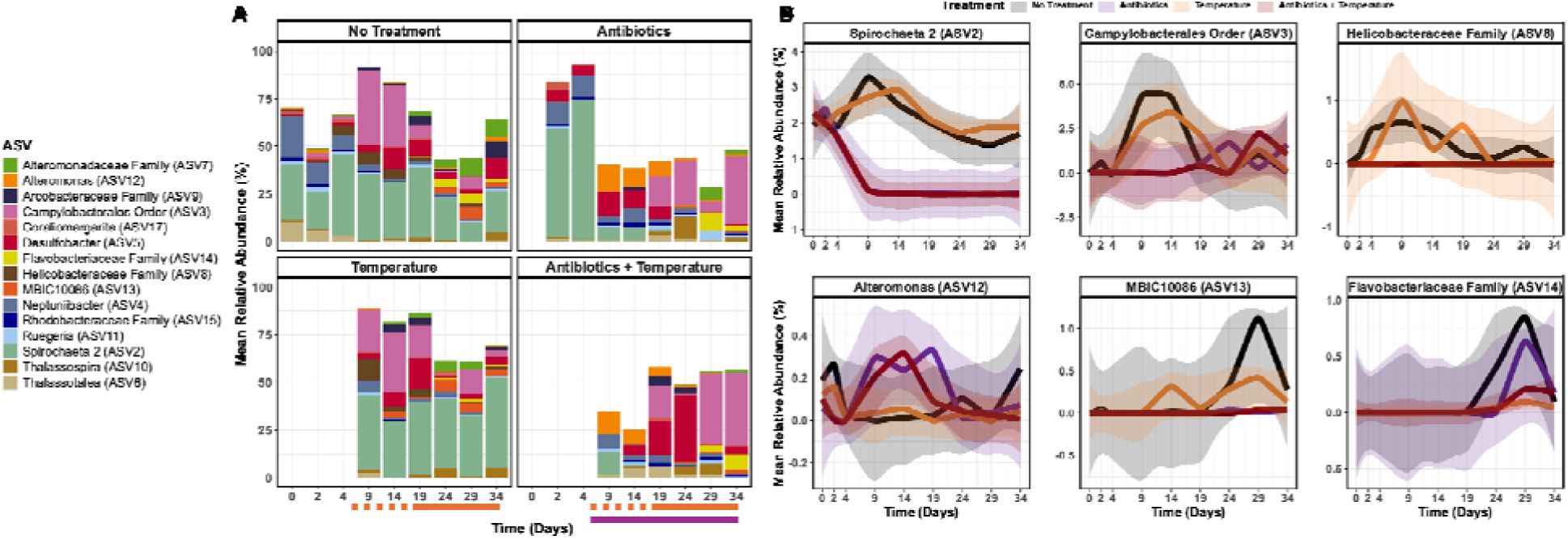
A) Relative abundance of the top 15 most abundant ASVs after *Ca.* Aquirickettsia rohweri was removed. B) Relative abundance over time for individual ASVs. Ribbons represent the 95% confidence interval.

## Discussion

Using both 16S relative abundance data and qPCR methods, our results indicate that *Ca.* Aquirickettsia rohweri is able to maintain microbiome dominance despite antibiotic challenge, sub-bleaching thermal stress, and a sequential disturbance of the two. However, while *Ca.* Aquirickettsia rohweri was not affected, our treatments did affect minor taxa, resulting in sustained shifts in microbiome composition and diversity within the rare biosphere.

### Sustained effects of antibiotic challenge on coral microbiomes

Some acroporid species are thought to maintain relatively stable microbiomes that may recover rapidly following disturbances – likely owing to strong host influence on microbiome structure (Aguirre et al., 2022; Dunphy et al., 2019, 2021; Glasl et al., 2019). Stability, resilience, and recovery, however, refer to suite of responses such as maintenance of major bacterial taxa or core community structure, or maintaining bacterial functional guilds (Shade et al., 2012), and conclusions are often species- or body site-dependent (Lima et al., 2020; Marchioro et al., 2020). In the present study, we found that the dominant microbiome fraction remained largely compositionally stable across all treatment groups, as a single *Ca.* Aquirickettsia rohweri ASV constituted upwards of 99% of the disease-susceptible *A. cervicornis* microbiome, while significant changes in the minor fraction resulted in sustained shifts in microbiome composition and diversity. Consistent with our findings, a recent study found that the dominant taxon within Panamanian *A. cervicornis* microbiomes, *Endozoicomonas*, remained in stable association with the host after antibiotic challenge and subsequent field transplantation to a common garden (Dunphy et al., 2021). Results from the present study, however, clearly indicate that although there is a stable association of the dominant taxon after antibiotic challenge, changes in minor taxa drove significant differences in both diversity and composition that persisted over a three-week recovery period. One limitation in our study, however, is that samples were not taken beyond three weeks, and thus whether or not these microbial communities remain significantly disrupted by antibiotics in the long term.

Shorter-term studies have found that antibiotic-disrupted Acropora muricata microbiomes begin to recover 12 hours after antibiotic cessation (Sweet et al., 2011b) and Astrangia poculata mucus microbiomes return to similar pre-disturbance states after a two-week recovery period (Bent et al., 2021). These results, however, are likely dependent on coral species, body site sampled, as well as the presence of future disturbances. Furthermore, in addition to being a different species than in the present study, mucus microbiomes may also display more rapid recovery after disturbance given its role as a first line of defense against disturbances, and mucus composition is thought to correlate more with environmental factors (Lima et al., 2020; Marchioro et al., 2020) whereas tissue microbiome composition is often correlated with host factors (Marchioro et al., 2020). Therefore, the observed difference in sustained disturbance versus rapid recovery is likely due, in part, to the overwhelming dominance of *Ca.* Aquirickettsia rohweri. In addition to the parasitic nature of *Ca.* A. rohweri, microbiome dominance itself is thought to present ecological tradeoffs that may be only circumstantially beneficial (Epstein et al., 2025). Therefore, despite *Ca.* A. rohweri-dominant *A. cervicornis* appearing outwardly healthy, the effects of this dominance in the face of disturbance may manifest as a reduced capacity for its microbiome to recover to pre-disturbance composition – further warranting deeper investigations into how antibiotics affect microbiome recovery trajectory of both diseased and non-diseased corals.

### Effects of antibiotics on the rare microbial biosphere

Antibiotics, sub-bleaching thermal stress, and a sequential combination of the two each contributed to significant changes in relative abundances of minor microbiome members. Most notably, *Spirochaeta 2* (ASV2) had significantly lower abundance in the antibiotic-treated samples compared to the control and temperature only samples. *Spirochaeta* ASVs are often found in Floridian *A. cervicornis* (Gignoux-Wolfsohn et al., 2020; Klinges et al., 2022, 2023; Williams et al., 2022), and in other coral species throughout the Caribbean (Arriaga-Piñón et al., 2024; Pratte & Richardson, 2018; Rosales et al., 2020). Spirochaetaceae are common among corals, yet their roles remain largely unknown. However, taxa within this family are hypothesized to play roles in nitrogen cycling in soft corals (Park et al., 2022), and their relative abundances may correlate with *A. cervicornis* survivorship under elevated nutrients (Palacio-Castro et al., 2022). Therefore, the sustained reduction in *Spirochaeta 2* relative abundance observed in the present study may have negative impacts on host response and recovery to future disturbances such as nutrient enrichment – although further research is required. Furthermore, several studies have found *Spirochaeta 2* in both diseased and healthy *A. cervicornis* and *A. palmata*, thereby complicating the potential relationship with the coral host (Gignoux-Wolfsohn et al., 2020; Rosales et al., 2019; Young et al., 2023). Young et al., hypothesize that certain *Spirochaeta 2* species may be present in apparently healthy coral microbiomes, while other species are present in pathobiome states (Young et al., 2023). Provided that many studies use short-read sequencing, functional inference and finer-resolution taxonomic information is often limited; therefore, future efforts utilizing full-length 16S rRNA gene sequencing may uncover species or strain-level resolution in order to inform this hypothesis.

Although the *Campylobacterales* ASV3 was not deemed differentially abundant by MaAsLin2, its relative abundance patterns between treatments suggests that it is likely susceptible to antibiotics, but may increase in relative abundance to levels comparable to ‘no treatment’ and ‘temperature’ groups after a recovery period of about two weeks. The perceived increase in relative abundance of this ASV in ‘no treatment’ and ‘temperature’ samples likely coincides with the change from enclosed to flow-through aquaria in which fresh seawater is constantly circulating in the system and likely enriching for *Campylobacterales*. We previously found that this same *Campylobacterales* ASV, present in high relative abundance in disease-resistant *A. cervicornis*, is highly susceptible to antibiotics (Patton et al., 2025). Yet due to the time constraints of that study, we were unable to determine whether or not this ASV would recover to pre-disturbance levels. Although genotypic differences likely drive microbiome re-assembly, our results indicate that after temporary suppression *Campylobacterales* may effectively recover after antibiotic disruption.

After a recovery period, antibiotic-challenged samples displayed a significant reduction in *Caedibacter taeniospiralis* (ASV26) relative abundance (100% BLASTn sequence identity to *Cysteiniphilum litorale* isolated from seawater). Along with *Vibrio* sp., *Cysteiniphilum litorale* has been recently implicated as a putative pathogen in White Band Disease (Selwyn et al., 2024, 2025; Trytten et al., 2025). Interestingly, *C. litorale* was found to increase in relative abundance in antibiotic-treated, disease-resistant *A. cervicornis* as a potential opportunistic pathogen (Patton et al., 2025), yet decreased in relative abundance in the disease-susceptible *A. cervicornis* in the present study. This suggests that antibiotics may enrich for opportunistic establishment in disease-resistant *A. cervicornis,* yet reduce this potentially pathogenic bacterium in a disease-susceptible genotype, which further emphasizes the importance of understanding how antibiotics differentially affect non-diseased conspecifics, as well as those with variable degrees of disease-susceptibility. However, it must be noted that these were not identical ASVs and only shared 97.5% sequence identity. Selwyn *et al*, found that prophylactic treatment of healthy *A. cervicornis* with kanamycin, ampicillin, chloramphenicol, and tetracycline significantly reduced infection rates when exposed to WBD, and *C. litorale* was significantly associated with disease outcome (Selwyn et al., 2025). Given that Panamanian *A. cervicornis* used in the Selwyn *et al*. study had microbiomes dominated by *Endozoicomonas* rather than *A. cervicornis*, future research efforts should assess how prophylactic antibiotic administration affects WBD transmission in *Ca.* A. rohweri-dominant *A. cervicornis*.

### Effects of temperature and subsequent stressor challenge on rare microbial biosphere and *Ca.* Aquirickettsia rohweri

Temperature challenge revealed only subtle differences in microbiome diversity, composition and differential abundance – likely due to a combination of the gradual temperature ramp pace and the constraints on the total temperature increase. No differentially abundant taxa were uniquely associated with the ‘temperature’ treatment group. During the temperature challenge phase in the ‘antibiotics + temperature’ group, we observed two potential opportunistic ASVs that increased in relative abundance, yet did not appear differentially abundant in the ‘antibiotics’ or ‘temperature’ treatment groups alone. Cyanobacteria Class (ASV182) had significantly higher relative abundance in the ‘antibiotics + temperature’ group compared to the ‘no treatment’ group during the temperature challenge phase, and shares 96% BLASTn sequence identity with an unclassified Cyanobacteria isolated from a Caribbean sponge, *Xestospondia muta*, infected with sponge orange band disease (Angermeier et al., 2011). Cyanobacteria have long been implicated in coral disease – specifically in the cyanobacterial mats associated with Black Band Disease (BBD) (Frias-Lopez et al., 2004). *Thalassotalea* (ASV6) is another potentially opportunistic ASV with significantly higher relative abundance in the ‘antibiotics + temperature’ samples during the temperature challenge phase. This ASV shares 100% BLASTn sequence identity with an unclassified bacterium isolated from a BBD microbial mat affecting the coral *Colpophyllia natans* (Klaus et al., 2011). Several ASVs within this genus were also identified as putative WBD-associated opportunists (Trytten et al., 2025). Despite antibiotics reducing the relative abundance of a purported WBD pathogen, a synergistic effect is observed in which the combined stressors increase the relative abundance of two potentially pathogenic bacteria that were not deemed differentially abundant in either of the two stressors alone.

Differential abundance analysis revealed that *Ca.* A. rohweri significantly increased in relative abundance, yet past research has shown that *Ca.* Aquirickettsia rohweri is lost during thermally-induced bleaching (Klinges et al., 2020). Therefore, we hypothesized that even at elevated temperatures below the bleaching threshold we may see a reduction in *Ca.* A. rohweri. However, during the temperature challenge phase, we saw that *Ca.* A. rohweri relative abundance significantly increased in all treatment groups compared to the control. Bioenergetic models suggest that moderate temperature stress increases corals’ metabolic rates (Pfab et al., 2024), which may explain, in part, the perceived increase in *Ca.* A. rohweri relative abundance. When the timepoints were separated and relative quantification was performed, only a single timepoint in each of the treatment groups displayed significantly higher abundance *Ca*. A. rohweri compared to the control samples, while the general trend in both the ‘temperature’ and ‘antibiotics + temperature’ groups was a decrease in *Ca.* A. rohweri abundance over time. Therefore, our results may indicate that, up to a certain, currently undefined, threshold, increased temperatures may support *Ca.* A. rohweri proliferation as it acquires additional host metabolites. Yet after the threshold is surpassed, even at sub-bleaching temperatures, temperature stress becomes detrimental to *Ca.* A. rohweri survival.

### *Ca.* Aquirickettsia rohweri intracellularity and microbiome dominance

Bacterial intracellularity imparts a degree of microbiome compositional stability in the face of antibiotic disturbance due to the physical protection afforded by host cells. Although some classes of antibiotics used in this study can cross eukaryotic membranes via phagocytosis or diffusion, they may remain ineffective once inside the host cell due to mechanisms such as bacterial efflux pumps reducing antibiotic bioaccumulation and ultimately efficacy (Webber & Piddock, 2003). Genomic information reveals that *Ca.* Aquirickettsia rohweri is capable of sensing and responding to extracellular stimuli via coupled ATP-binding cassette transporters and two-component systems (Klinges et al., 2019). Furthermore, the *Ca.* A. rohweri genome also contains genes necessary for a type IV secretion system (Klinges et al., 2019) which, in addition to a suite of other functions, has been implicated in the spread of antimicrobial resistance (Cascales & Christie, 2003). Although the *Ca.* A. rohweri genome contains the molecular machinery necessary for active antibiotic resistance, it is unknown whether or not they are expressed. Therefore, future efforts aimed at isolating *Ca.* A. rohweri in culture would help inform whether there are intrinsic genomic features allowing for antibiotic resistance or resilience, or if host-association is what maintains the stable association. Understanding this would also contribute to prophylactic intervention to prevent *Ca.* A. rohweri microbiome dominance.

## Conclusion

In this study, we found that the dominant, putatively parasitic bacterium *Candidatus* Aquirickettsia rohweri was not negatively affected by a suite of broad-spectrum antibiotics, sub-bleaching thermal stress challenge, or a subsequent disturbance of antibiotics and sub-bleaching thermal stress in a disease-susceptible *Acropora cervicornis* genotype. Although prophylactic and retroactive antibiotic treatments have effectively treated and prevented some diseases in scleractinian corals, antibiotics may not be a viable treatment for *A. cervicornis* genotypes whose microbiomes are dominated by *Ca.* A. rohweri, as this bacterium was found to proliferate in response to antibiotic administration. Furthermore, given that this bacterium cannot be treated with antibiotics, it is especially important to understand *Ca.* A. rohweri transmission and persistence dynamics in order to develop microbiome intervention strategies for restoration success.

## Acknowledgements

We would like to acknowledge staff at Mote Marine Laboratory’s IC2R3 for assisting with specimen collection and CAOS system monitoring. The authors thank Drs. Thomas Sharpton, Ryan Mueller, Maude David, and Xiaoli Fern for their feedback on the formal analysis. We would also like to thank the Center for Quantitative Life Sciences RRID:SCR_018373 at Oregon State University for the sequencing services that generated data for this publication. We also acknowledge the use of the Biological Nanostructures Laboratory within the California NanoSystems Institute, which is supported by the University of California, Santa Barbara and the University of California, Office of the President. We would also like to thank the National Science Foundation for supporting this work.

## Data Availability

Scripts used for bioinformatic and statistical analysis can be found at: https://github.com/pattonsunni/RoL_AntibioticsTemp_G50. The 16S rRNA dataset supporting the conclusions of this article is available in the NCBI Sequence Read Archive (SRA) repository under the BioProject accession number PRJNA1489297.

## Contributions

S.P. conceptualized the study, performed the investigation, conducted the formal analysis, generated all figures, wrote the original draft of the manuscript, reviewed and edited the manuscript, and acquired funding. E.F. conceptualized the study, performed the investigation, and reviewed and edited the manuscript. L.S. performed the investigation and reviewed and edited the manuscript. G.K. Reviewed and edited the manuscript. E.M.M. Acquired funding and reviewed and edited the manuscript. R.L.V.T. conceptualized the study, performed the investigation, acquired funding, provided supervision, wrote the original draft of the manuscript, and reviewed and edited the manuscript.

## Supporting Information

**Figure S1.**
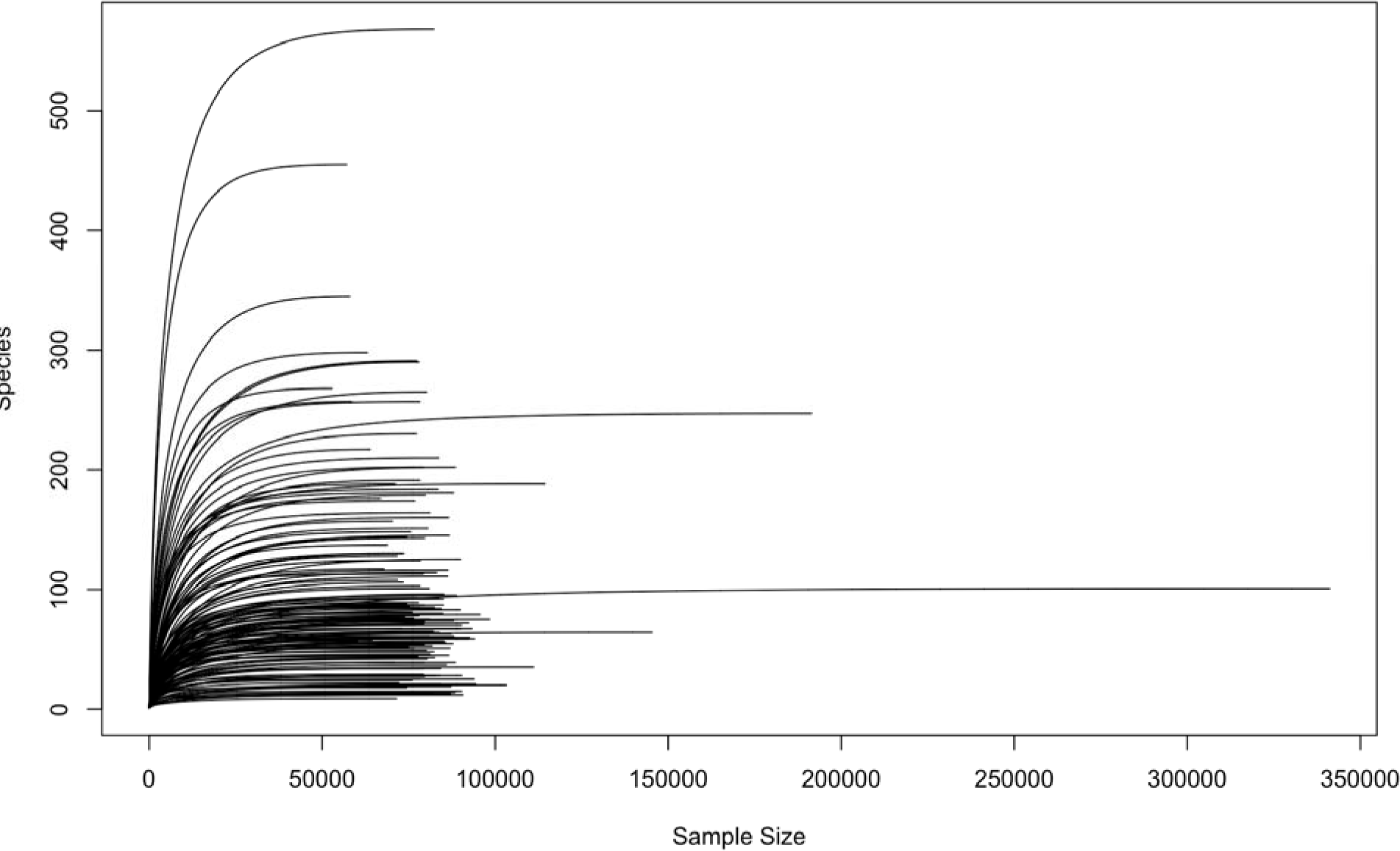
Rarefaction curve displaying the library size that adequately captures the species richness for each sample. Phyloseq object was rarefied to 52,706 reads which captures a majority of the species richness without losing any samples.

**Figure S2.**
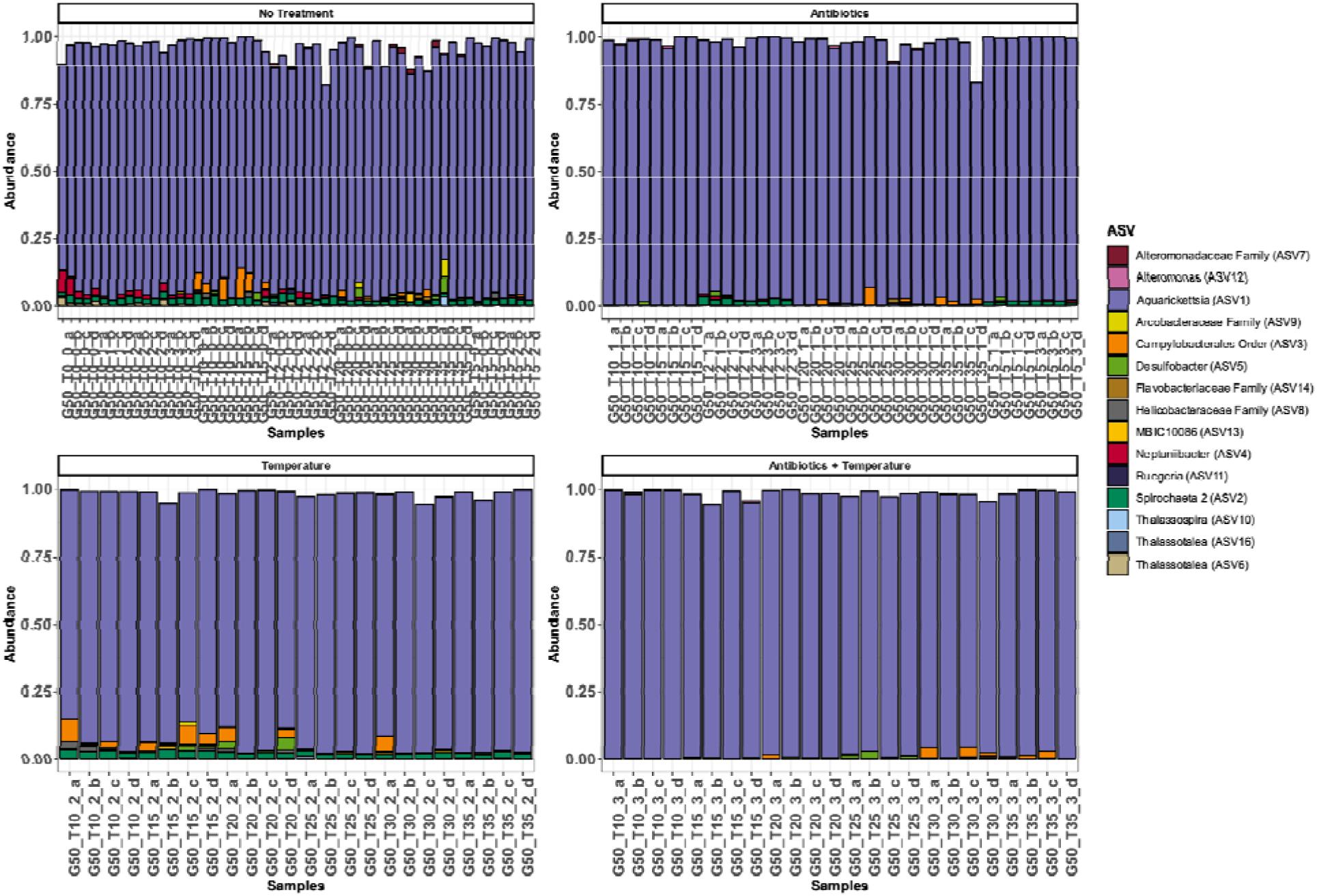
Relative abundance plot of the top 15 most abundant ASVs. Bars represent individual coral samples and the plot is faceted by treatment group.

